# Benefits and costs of a global cooperative surveillance strategy to control trans-boundary crop pathogens

**DOI:** 10.1101/2022.10.13.512036

**Authors:** Andrea Radici, Davide Martinetti, Daniele Bevacqua

## Abstract

Trans-boundary diseases are extremely complex to control and can cause global socioeconomic damages. In the context of crop protection, surveillance strategies are usually designed according to administrative boundaries. In this study, we investigate to which extent this geographical scale of surveillance is suitable for long-distance dispersed pathogens. We lever on a global epidemic network, presented in a previous work, describing worldwide potential transport of *P. graminis*, the causal agent of stem rust of wheat. We conceive two surveillance strategies: “Country-based” and “Cooperative” and we compare their performances, in terms of minimizing the number of sentinels deployed, to achieve given surveillance targets, both at the global and country level. As expected, we find that a “Cooperative” strategy is more efficient at the global scale, and this is particularly true for intermediate targets of surveillance. However, costs and benefits of adopting a “Cooperative” strategy are not equally distributed among countries. Medium size countries in central Europe and Asia are those that would benefit most from a cooperative strategy. On the other hand, Greece and Finland, having a small wheat production but placed in important *Puccinia* pathways, are those that should deploy, in the global interest, more sentinels than they would place in the national interest. Among the major wheat producers, China is the only country that would have a cost from a cooperative strategy, while India, Russia, United States, France and Ukraine would have the most benefits. It follows that the acknowledgement of these discrepancies could help to achieve general stakeholder support for a global international cooperative surveillance system.

## 1. Introduction

The issue of surveillance of trans-boundary diseases has recently came in the spotlight due to the Covid-19 pandemic [1, 2, 3]. Trans-boundary diseases are highly contagious, epidemic diseases, with the potential to spread rapidly across the globe and causing substantial socioeconomic and health issues. First outbreaks of such diseases [4, 5], as well as biological invasions of alien species [6], are hardly predictable events. Furthermore, lack, mismatch or delay in the communication of first detections among countries, together with uncoordinated control measures, may lead to inefficient management [7, 8]. Notably, the threat posed by airborne crop pathogens represents a paradigmatic case of trans-boundary spread [9, 10, 11]: the risk of large losses in food production due to unexpected outbreaks has prompted researchers and institutions to explore international surveillance systems to timely tackle the diffusion of the most alarming crop pathogens [12, 7]. The spatio-temporal persistence of seasonal movements, such as the well known *Puccinia* pathway from Mexico to Canada [13, 14], inspired recent studies to conceive innovative surveillance systems [15, 16, 17, 18]. In spite of such efforts, standard surveillance of trans-boundary crop diseases is still mainly performed according to administrative boundaries, regardless of the actual scale of spread of the disease, often lacking international and timely communication of first detections [12, 7, 19]. Yet, the benefit of a global, cooperative and communicative strategy [20] over a country-based one, remains unknown.

In this study, we investigate to what extent, and under which conditions, administrative boundaries represent a suitable scale for surveillance of long-distance dispersed crop pathogens, and whether international cooperation would make crop protection more effective. We considered stem rust of wheat, caused by fungal pathogen P. *graminis*, as a case of study. Having reappeared in Western Europe after several decades of absence [21, 11, 5], this pathogen is considered a threat to global food security due to the rapid spread of virulent races through a globally distributed host [22, 19]. In a recent article, we retraced its global epidemic network across worldwide wheat-producing countries [18]. In the present study, we lever on this epidemic network to conceive two surveillance strategies, a “Country-based” one, representing a within-boundary scenario with no collaboration and communication between countries, and a “Cooperative” one where countries look after each others territories and timely communicate disease detections. We compare their performances in terms of surveillance effort needed to achieve given targets both at the global and local scale. We found that a “Cooperative” strategy is generally convenient at the global scale, especially at intermediate targets of surveillance, but costs and benefits are not homogeneously shared among countries.

## 2. Materials and methods

### 2.1. Case study

*Puccinia graminis* f. sp. *tritici* is an airborne fungal pathogen responsible for the stem rust of wheat. Its spores can be transported over long distances by wind at continental scales [23] and may cause severe losses to its main host, wheat, which represent the most diffuse agricultural land cover. Every year, more than 750 × 10^6^ metric tons of wheat are produced worldwide, with countries such as China (133.6), India (103.6), Russia (74.5), United States (52.3) and France (40.6) accounting for more than 50% of the total production [24]. In the majority of these countries the presence of *P. graminis* has been controlled by the use of resistant cultivars and the eradication of its secondary host, *B. vulgaris*, which enables overwintering in temperate regions, while recent reappearances have been mitigated by fungicide use [25, 26, 21, 27, 28].

### 2.2. *The worldwide* Puccinia *epidemic network*

In order to evaluate the performances of different surveillance strategies, we levered on the epidemic networks obtained in a previous study, see [18] for details. In the following paragraphs we present a summary of the methodology proposed there.

In Radici *et al*. [18], we simulated worldwide transport of *P. graminis* spores among wheat producing countries, obtaining a time-varying connectivity network **C**. In **C**, the 7,814 nodes represent 0.5° × 0.5° cells (≈ 2,000 km^2^) of wheat-producing countries [29], while edges represent air-mass connections among cells, computed at a time resolution of 6 hours for the time span 2013-2018. More specifically, edges are computed in such a way that they account for the likelihood of air-mass trajectories (computed via NOAA’s HYSPLIT model [30]) connecting a release node (where the host is available and environmental conditions are favourable for sporulation) and an arrival node (where the host is available and environmental conditions are favourable for infection). Trajectories are filtered with a number of criteria (rain washout, cumulative UV radiation, flight duration and altitude) to exclude those air-mass movements that are less likely to lead to an effective spore transport event.

### 2.3. Surveillance strategy design

We further considered the problem of establishing a reduced set of *sentinels*, nodes where the presence of the pathogen is systematically monitored (i.e., the surveillance effort), that should guarantee the largest aggregated coverage of the domain (i.e., the surveillance target) and provide an early-warning system for the appearance of the pathogens [18]. First of all, we defined the *coverage* of a sentinel as the set of nodes that points towards it, under the assumption that, by monitoring the presence of the pathogen in a sentinel, we can indirectly observe the possible presence in all those nodes that are pointing to it. We levered on an iterative heuristic algorithm to determine sub-optimal solutions to the problem of finding the smallest set of sentinels that guarantees the maximum aggregated coverage. It consist in: 1) finding the node associated to the largest coverage, 2) add this node to the sentinel set s_*σ*_, initially empty, 3) label its coverage as surveilled and remove all the edges outgoing from the surveilled set, 4) repeat steps 1-3) until the proportion of nodes in the aggregated coverage reaches the desired target *σ*. The optimal set of sentinels s_*σ*_ is ranked by growing aggregated coverage. The size of s_σ_ defines the surveillance effort *x_σ_*.

### 2.4. Strategy design: “Cooperative” vs “Country-based”

As explained in the previous section, the iterative heuristic algorithm allows to find the minimum sentinel set size for a desired target *σ* at the global scale. While this is straightforward from a computational point of view, it does not represent how real surveillance works, since surveillance may be more likely conducted by each country (or even region) independently. In fact, this algorithm assumes that sentinel locations are chosen regardless of administrative borders, while it may not be the case. For these reasons, we named the solution of the above-mentioned algorithm as the “Cooperative” strategy.

Alternatively, we built a second strategy where surveillance is designed mimicking a more realistic scenario. This strategy, named “Country-based”, differs from the previous as sentinel prioritization is carried out each country independently of the others, which means that the set cover algorithm is solved at the country level (Fig. 1). To facilitate this operation, we assumed that the coverage of a sentinel is limited by the administrative borders of the country in which it is located. This is equivalent to modify the connectivity network **C** by labelling each node with the name of the country where it is placed and removing all border-crossing edges, thus obtaining a new network **C**^−T^. Then, we run the same iterative heuristic algorithm described in the previous paragraph, independently for each country, using the network **C**^−T^ to obtain the optimal sentinel sets 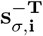 for each country *i*, ranked by growing aggregated country coverage. To compare the global performances of the “Cooperative” and “Country-based” strategies, we computed the number of sentinels needed to achieve different global targets.

**Figure 1:**
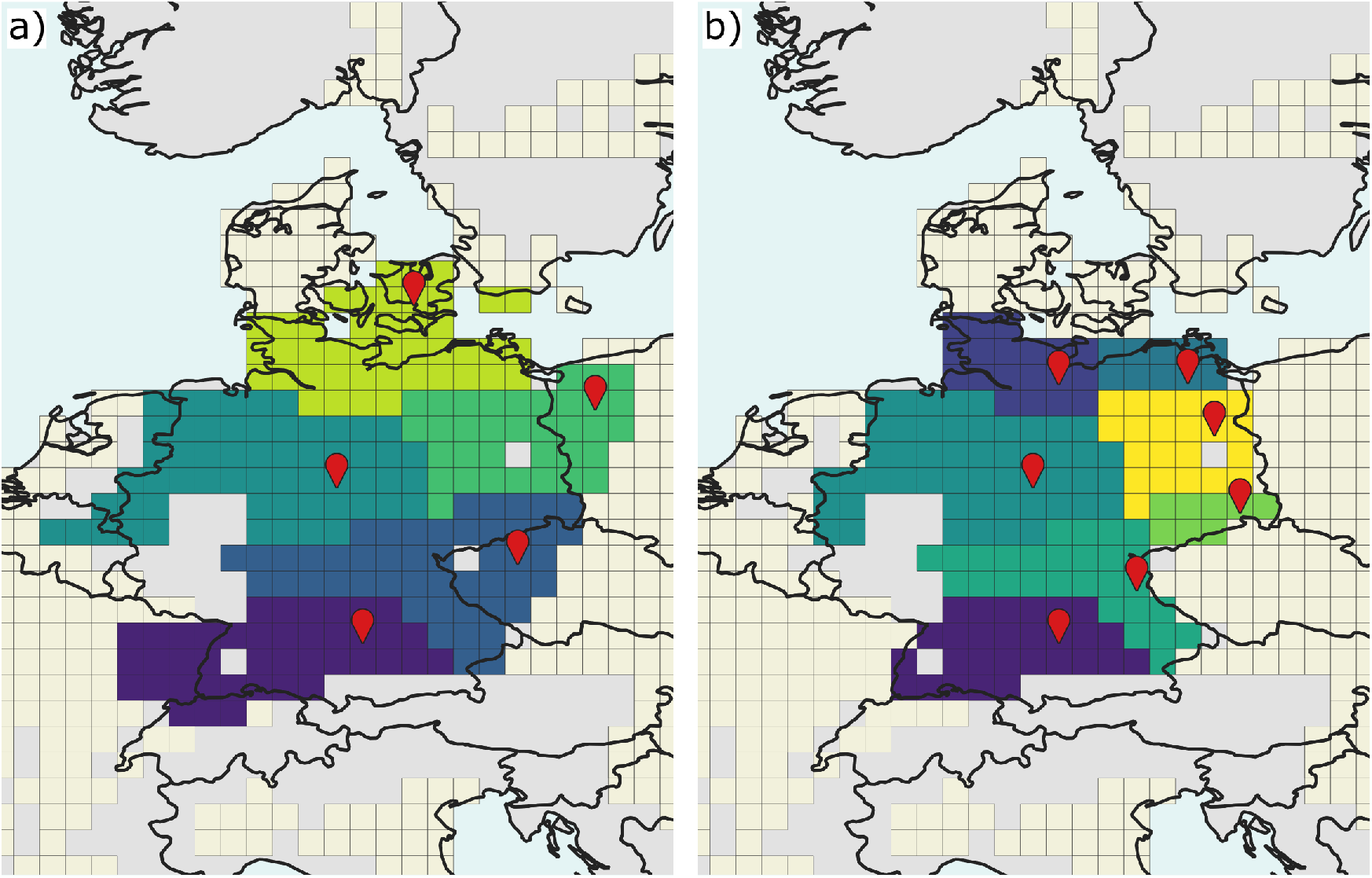
A graphic example to compare “Cooperative” (a) and “Country-based” (b) strategies. Pale yellow squares represent nodes of the networks, corresponding to wheat production regions. In the “Cooperative” strategy, five sentinels are needed to cover all nodes in Germany: three of them being located abroad (Denmark, Poland, Czechia), while those located in Germany also contribute to cover cells in the Netherlands, France and Switzerland. In the “Country-based” strategy, seven sentinels, all placed in Germany, are needed to surveil German nodes while they do not contribute to trans-boundary surveillance.

### 2.5. Benefits and costs of cooperation at country scale

To investigate how the burden of surveillance is shared among countries, for each country *i*, we calculated the number of sentinels *x_i,σ,s_* needed to achieve a country target of *σ* under a given strategy *s* (*s* = “Cooperative” or “Country-based”). Then, we defined the index *α_i,σ_* as follows (Eq. 1):

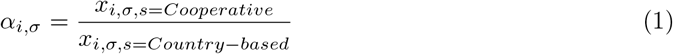

For a given target *σ, α_i,σ_* measures the ratio between the number of local sentinels needed to achieve *σ* in the “Cooperative” and in the “Country-based” strategy, for a given country *i*. The interpretation of *α_i,σ_* is rather straightforward: if *α_i,σ_* < 1, country i requires less sentinels within its borders in the “Cooperative” scenario than in the “Country-based” one for achieving the same surveillance target *σ*. If *α_i,σ_* > 1, the opposite is true, while if *α_i,σ_* = 1, country *i* needs the same number of sentinels in both the strategies for achieving surveillance target *σ*. We evaluated *α_i,σ_* for *σ* = 1%, 2%, … 100% and then we calculated mean 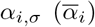 by country. Depending on the values of 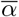, we classified each country either as *i*) “CoopBeneficial” if, on average, less sentinels are needed with the “Cooperative” strategy to achieve the same target 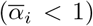, ii) “CoopNeutral” if the number of sentinel is the same 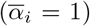 or eventually iii) “CoopAdverse” more sentinels are needed in the “Cooperative” scenario 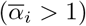.

After having computed 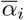 by country, we aggregated it by continent to investigate geographical heterogeneity of benefits and costs of cooperative surveillance.

### 2.6. Robustness of the sentinel sets

To assess the temporal robustness of the performances sentinel sets to slight changes in the epidemic network, we set up a validation procedure consisting in projecting the yearly epidemic networks of [18] into:

1. a design (directed, binary) network **C**_D_, generated by considering recurring connections occurring at least once a year and at least three times over the 4-year interval 2013-2016 (i.e., ≥ 75% of the years);
2. a validation (directed, binary) network **C_V_**, obtained by considering only those connections occurring at least once a year both in 2017 and 2018.

We then used the network **C_D_** to design the surveillance strategy (i.e., identify the sentinel sets 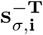 and **s**_*σ*_), while we used network **C_V_** to assess its robustness to temporal changes of connectivity patterns.

## 3. Results

### 3.1. Surveillance effort reduction due to cooperation depends on the desired coverage level

Under the “Country-based” strategy, an aggregated coverage of the worldwide wheat producing regions (*σ* = 100%) can be achieved by monitoring 1,148 sentinels, i.e. 14.7% of the total number of nodes (Fig. 2a). A “Cooperative” strategy allows to slightly reduce the sentinel set size to 1,007 sentinels for a complete coverage (12.9%, equivalent to a reduction of −12% with respect to the “Country-based” strategy, Fig. 2b). However, the advantage of cooperation is more evident for intermediate values of *σ*. For example, monitoring at least *σ_i_* = 50% of the wheat producing regions by country in a “Country-based” strategy would require 209 sentinels (2.7% of the nodes). Due to the discrete nature of each coverage, this would correspond to a worldwide target of about *σ* = 58% (Fig. 2a). With 209 sentinels, the “Cooperative” strategy would achieve a worldwide coverage of *σ* = 78%. Similarly, the same target of *σ* = 50% would ask for 64 sentinels under the “Cooperative” strategy (i.e. 0.8% of the nodes, or a reduction of 69% with respect to the “Country-based” strategy), while an aggregated coverage of 58% would be obtained with 87 sentinels (1.1%, or −58% with respect to the “Country-based” strategy; Fig. 2b).

**Figure 2:**
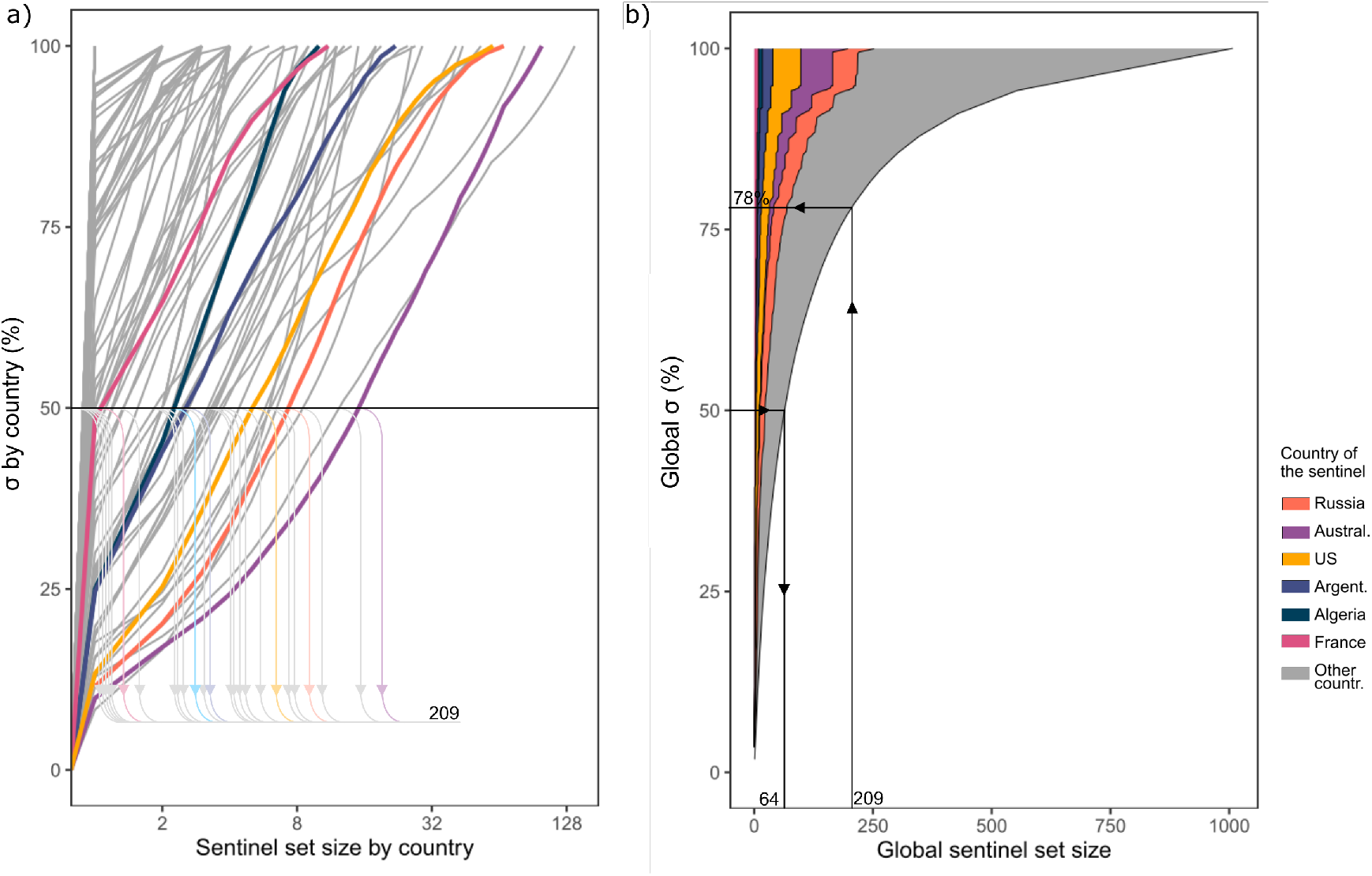
Panel a) shows the number of sentinel nodes needed in each country to achieve increasing surveillance targets *σ* in the “Country-based” strategy. One country for each continent is highlighted as example. Notice that the horizontal axis of this panel is in logarithmic scale. By summing the number of sentinels for each line/country when they intersect the horizontal line corresponding to *σ* = 50%, we obtain a total of 209 sentinels needed to cover at least 50% in each country. Panel b) shows the number of sentinel nodes needed in the “Cooperative” strategy to achieve increasing worldwide target *σ*. We can observe that 209 sentinels would allow a coverage of 78%, while the target *σ* = 50% may be achieved with just 64 nodes.

### 3.2. Heterogeneity in the distribution of surveillance effort reduction due to cooperation

Overall, out of 87 countries, 55 (63%) are classified as CoopBeneficial, 23 (27%) as CoopNeutral, and 9 (10%) as CoopAdverse. In terms of wheat production, around 71% is located in CoopBeneficial countries, 6% in CoopNeutral countries while 23% in CoopAdverse ones (Fig. 3a).

**Figure 3:**
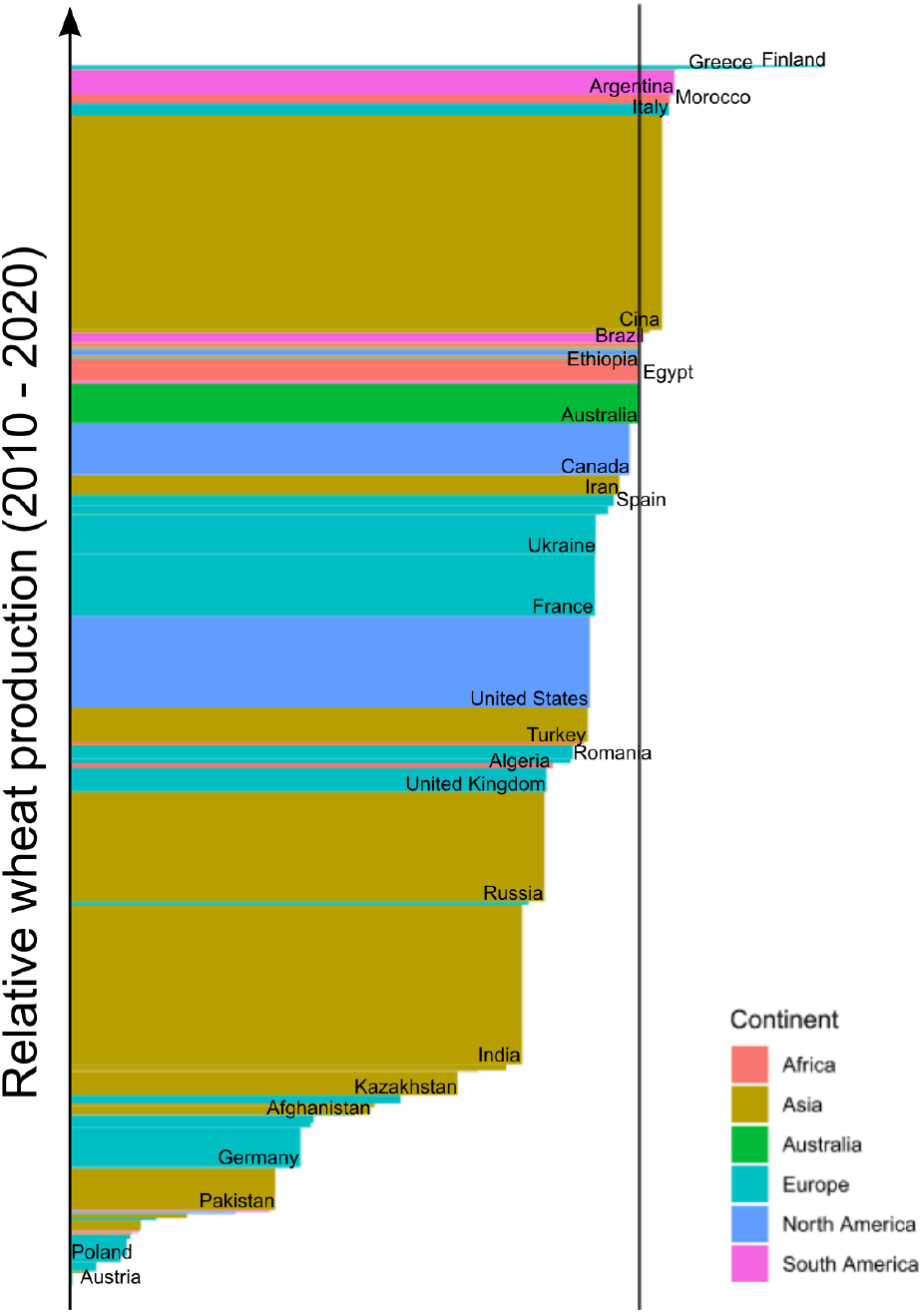
Barplot of the index 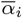 for all wheat-producing countries considered in the study. Each country is represented by a rectangle where the base is proportional to 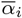 and the height to wheat production in 2010 - 2020 according to [24]. 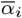 is calculated against the design networks **C_D_**.

We also assessed the difference in the distribution of the surveillance effort under the two strategies by differentiating large (at least 45 nodes), medium (between 44 and 13 nodes) and small countries (12 or less nodes; Fig. 3 and SI1). This naming reflects the wheat surface, and does not refer to the true country size. For 47 countries, mainly medium (e.g., Czechia or Uruguay) or large (e.g., India or Russia), *α_i,σ_* is always ≤ 1, thus implying an advantage in adopting a “Cooperative” strategy independently of *σ*. Only 4 countries (Morocco, Greece, Finland and Nepal) are always discouraged from adopting a “Cooperative” strategy. Great part of the small countries (such as Yemen or New Zealand) display *α_i,σ_* = 1 ∀*σ*, for which the two strategies are equivalent. For a few number of large (e.g. US, China or Iran) or medium countries (e.g. Moldova or Tunisia), *α_i,σ_* is lower or larger than 1 depending on *σ*.

At the world scale, each continent (except Australia) has at least one CoopBeneficial, one CoopNeutral and one CoopAdverse country (Fig. 4a). In North America, countries are typically CoopBeneficial, while South America is more balanced. Continental Europe is mainly CoopBeneficial, with some countries (Belgium, Luxembourg, Austria, Slovenia, Croatia, Bosnia and Herzegovina, Albania, North Macedonia) having 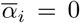. Finland has the highest 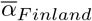 of 1.3. followed by Greece 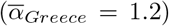. Asia has a composition similar to Europe, with few CoopAdverse countries (China, Mongolia, Nepal), some isolated CoopNeutral (e.g. Japan) and a majority of CoopBeneficial ones, mainly in inner parts of the continent. Africa is almost entirely CoopNeutral, with the exception of the Maghreb and Tanzania that are CoopBeneficial. Due to geographic isolation, island states such as Australia and New Zealand are CoopNeutral.

**Figure 4:**
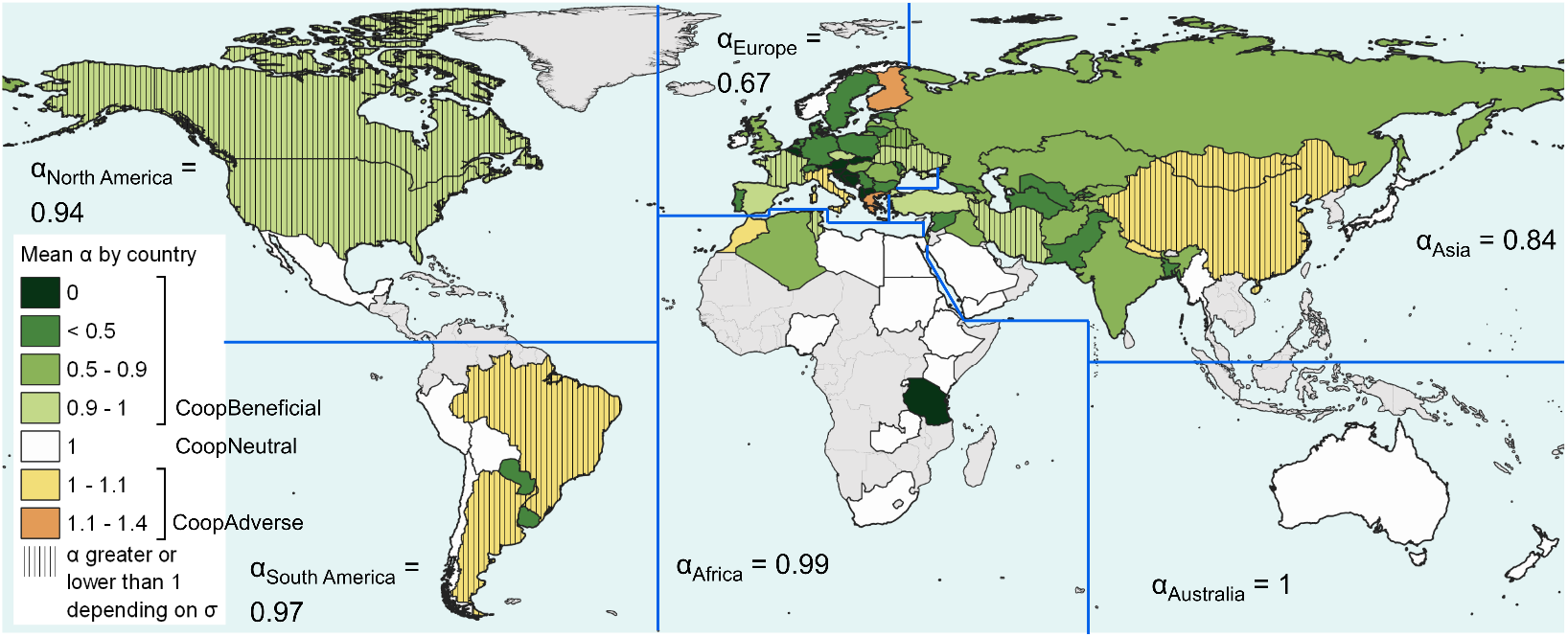
Global map of 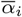. Surveillance performances are assessed against the design network **C_D_**. Average values by continents, weighted by country wheat production 2010 - 2020 [24] are also displayed.

### 3.3. Robustness of the surveillance strategies

We assessed the robustness of the performances of the sentinels sets prioritised via networks **C_D_** and 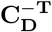 with the networks **C_V_** and 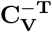. Overall, there is good agreement between the values of ā¿ using the design or the validation network for all countries *i*, as confirmed by a positive and statistically significant correlation coefficient of 0.89 (p-value < 2.2e — 16). A visual comparison is also provided in Fig.s SI2-5.

## 4. Discussion

### 4.1. From country-based to global cooperative crop protection

For whatever surveillance target *σ*, the “Cooperative” strategy is globally more efficient than a “Country-based” one. This difference is more significant at intermediate level of surveillance. These results have been corroborated by testing the sentinel sets against a new network **C_V_**, representing an air-masses connectivity which differs from those used in the design process. Despite the temporal variability of air masses movements, recurrent wind patterns, as well as temporal host availability and occurrence of environmental conditions favourable for infection, contribute to the seasonality of potential spore transport events. Amongst these connections, trans-boundary ones account for 23% of the total number of edges in the design network **C_D_** and 28% in the validation network **C_V_**. Our results suggest that they are relevant enough to affect the efficiency of a surveillance strategy.

As previous research have stressed, the scale of surveillance should correspond to that of the spread of the disease of interest, regardless of administrative boundaries [20]. We have collected evidence that, in the case of long-distance dispersed disease, a “Cooperative” approach allows to significantly reduce the surveillance effort needed to achieve a global coverage. This outcome agrees with previous studies, which underlined that neglecting long-distance connectivity leads to an underestimation of the disease spread capacity [31].

Despite increasing evidence of a global advantage in cooperative international surveillance, crop surveillance design is still mostly dictated by administrative boundaries rather than the actual scale of the pathogen spread [20, 7]. This mismatch between optimal and actual scale of action affects also other kinds of trans-boundary natural disasters, such as biological invasions by alien species. In this regard, Diagne *et al* [6] recently outlined that invasion-related economic damages are projected to increase in the next decades; one reason behind the inertia in the implementation of international and coordinated protection strategies may lie in the costs underestimation by the general public, stakeholders and decision-makers. This may be particularly true in the case of airborne diseases, either crop related or not, where the direct observation of such dispersal mechanism is virtually unfeasible [32, 21], and may discourage consideration by decision-makers.

### 4.2. Network-thinking in crop surveillance

The use of networks and network thinking to support crop protection strategies have been largely advocated in recent studies [31, 15, 33, 34, 35]. In the most simplistic way, these strategies rely on the identification of the nodes of the network that most contribute to spread the disease; or that, if vaccinated, reduce the disease size. Concerning surveillance, relevant nodes correspond to those that may allow early disease detection if systematically monitored [36]. Here, we used the set cover algorithm to prioritize nodes to be monitored, i.e., sentinels. Set cover algorithm iteratively selects the node associated with the highest coverage. In turn, coverage can be thought as a step-by-step updated version of the in-degree, i.e. the number of the edges pointing to a node, penalising those nodes whose coverage overlaps with that of nodes already labelled as sentinels. Other studies already noted that in-degree (or simply degree for undirected networks) is, as a general rule of thumb, a good proxy of both a good sentinel and a potential influential disease spreader [37, 38]. Moreover, in our work we proposed an hybrid network and geographical approach, in which metadata are associated to network components: each node is associated to the label of the corresponding country, and each edge is consequently labelled as “trans-boundary” or not. To our knowledge, this study is one of the first attempts to compare non-topological surveillance strategies, i.e. “Cooperative” and “Country-based”, and to quantify the heterogeneity in the allocation of the burden of “Cooperative” surveillance.

### 4.3. Sharing benefits and costs of cooperation

Form a global perspective, a “Cooperative” strategy is necessarily more efficient compared to a “Country-based” one, since it corresponds to an optimization subjected to fewer constraints. However, it is interesting to quantify how such strategy performs against a “Country-based” strategy at country level, since benefits and burden may not be equally shared.

To investigate this possibility, we propose the index *α_i,σ_*, that measures the ratio between the number of sentinels needed in a “Cooperative” strategy vs. a “Country-based” one. We observe that medium-sized countries located in an inner continental position, such as in Central Europe or Central Asia, are associated to lowest 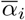 values, since they benefit of trans-boundary potential transport events among a landscape dominated by wheat producing areas. Insular countries, such as Australia, New Zealand or Japan, having no recurrent edges with other countries due to geographical isolation, are CoopNeutral. Due to the low presence of wheat, many African (excluded Maghreb) and South American countries are CoopNeutral. By contrast, it is more difficult to determine a rule of thumb for CoopAdverse countries. Finland and Nepal are small-medium sized wheat producing country, located at the point of arrival of western-eastern European [25] and Indian [13] “*Puccinia* pathways”, respectively. Given their relatively small size, they are forced to assume more sentinels in the benefits of upwind countries than they would need if let alone. By contrast, Canada, the final destination of the north-American pathway, is a large wheat producing country, hence it would need several sentinels no matter the strategy. We may suppose that, among those other countries which are CoopAdverse, Italy and Greece are strategically located in the middle of the Mediterranean basin and they may play as stepping stones [39] for epidemics spreading northward from Africa towards Central Europe; furthermore, both have relatively low wheat productions, hence they would need less sentinels if not cooperating, while Brazil, Argentina and China are large countries helping small and medium surrounding countries to decrease the size of their sentinel sets.

By averaging 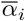 by continent it is possible to highlight those continents which would benefit the most of a cooperative surveillance. Europe and Asia display the lowest 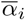 values (0.6-0.8), while for other continents it is generally around 1. To sum up, the connectivity network of this airborne disease creates a heterogeneous distribution of costs and benefits, but some continents would certainty take advantage of an international and cooperative surveillance system, namely Asia and Europe (Fig.s 4 and SI4).

The heterogeneous geographical distribution of impacts of cooperation suggests that a compensating mechanism should be set up to compensate local increases in costs. This idea can be borrowed from the socioeconomic concept of “burden sharing” [40, 41], which is finding application in the management of environmental goods. Differentiate GHG emissions reduction in the framework of the COP to achieve climate targets [42], as well as in the multi-stakeholders management of marine resources [43], may be two notably example. While our study tries to push towards a change in the perspective of governance of crop disease surveillance, we believe that proper identification of spatial distribution of costs and benefits can help facilitate international agreement for a global crop epidemic surveillance and gain support of all stakeholders.

## Supporting information

Supplementary material for the article

## 5. Acknowledgements

The authors acknowledge the support of funding from the French National Research Agency (ANR) for the BEYOND project (contract # 20-PCPA-0002) and SuMCrop Sustainable Management of Crop Health Program of INRAE that supported the work of all authors.

## Notes

### Competing Interest Statement

The authors have declared no competing interest.

