## Supplementary material for the article for "Benefits and costs of a global cooperative surveillance strategy to control trans-boundary crop pathogens"

### 1. Supplementary Methods and Results

#### 1.1. Local impacts of cooperation vs country size

In order to explore possible relations among surveillance effort distribution (i.e., CoopBeneficial, CoopNeutral and CoopAdverse) and country size, after having calculated $\alpha_{i,\sigma}$, we characterised each country by its wheat-producing surface: large (at least 45 cells), medium (between 44 and 13) and small (12 or less). Fig. SI1 reports a selection of the curves describing the relationship among country sentinel set size and aggregated coverage for some representative examples. The function $\alpha_{i,\sigma} = f(\sigma)$ is shown as well.

#### 1.2. Robustness of the surveillance strategies

We assessed the performances of same sentinels vector $\mathbf{s_i^{-T}}$ and $\mathbf{s}$ against the validation networks $\mathbf{C_V^{-T}}$ and $\mathbf{C_V}$ respectively. In both cases, the maximum achievable target $\sigma$ is lower than 100% (Fig.s SI2c,d) due to the reduction of the edges of the networks. However, it is still possible to estimate the sentinel set size needed to achieve a target of $\sigma_i = 50\%$ by country in the "Country-based" strategy, which is 459 (5.9% of the nodes; Fig. SI2c). This computation excludes Finland and Tunisia, for which the maximum aggregated coverage in $\mathbf{C_V^{-T}}$ are $\sigma_{Finland} = 25\%$ and $\sigma_{Tunisia} = 47\%$. Again, due to the discrete nature of the coverage, 459 sentinels are associated to a global coverage of 54%. In the "Cooperative" strategy, 459 sentinels would enable to increase the observed domain up to 75%, while the $x_{54}$ and $x_{50}$ are 138 (1.8% of the nodes) and 114 (1.5%) respectively, i.e. around a quarter of the sentinels needed in the "Country-based" strategy (Fig. SI2d).

*Corresponding author

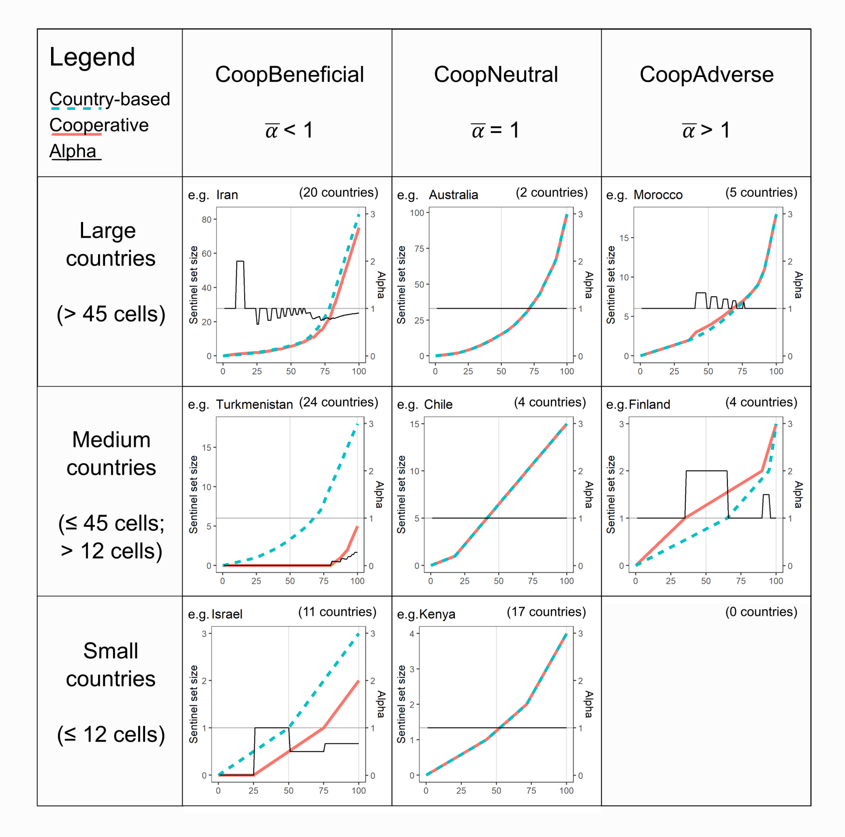


Figure SI1: Patterns of surveillance effort vs sentinel set size area investigated for different country size (rows) and values of $\overline{\alpha}$ (columns) for the two surveillance strategies. 63% of the countries is CoopBeneficial, and among them in 54% the "Cooperative" curve is always below the "Country-based" ($\alpha \leq 1$ $\forall \sigma$). For 27% of the countries, the two lines correspond, indicating no difference in terms of surveillance effort between the two strategies (central column). Eventually 10% are CoopAdverse (for 5% of the countries $\alpha \geq 1$ $\forall \sigma$). Few countries (15%) vary their $\alpha$ depending on $\sigma$ (Such as Iran, in the first row and first column). For each combination of country size and pattern of $\overline{\alpha}$, curves representing sentinel set size (first y axis) and $\alpha$ (second y axis) versus $\sigma$ (x axis) for a typical country are displayed, as well as the number of countries with a similar qualitative pattern.

*1.3. Robustness of the costs and benefit distribution among countries*

In a similar way compared to the previous paragraph, we assessed the robustness of the outcomes of the distribution of costs and benefits of a cooperative surveillance against the networks $\mathbf{C_V}$ and $\mathbf{C_V^{-T}}$. A visual comparison is given by observing Fig.s SI3 and SI4.

In terms of wheat production [1], passing from the design to the validating network implies that the fraction of CoopAdverse wheat producing countries slightly increases from 23% to 26%, CoopNeutral countries increase from 6% to 7% while the majority of the countries remain CoopBeneficial (from 71% to 66%, Fig. SI3a). We observe that 77 out of 87 countries (i.e., 89% of the countries) keep the same label as CoopBeneficial, CoopAdverse and CoopNeutral. The linear correlation coefficient in the scatterplot which compares $\overline{\alpha}_i$ computed with $\mathbf{C_D}$ and $\mathbf{C_V}$ is 0.89 (Fig. SI5), and the resulting p-value via the Pearson test is $< 2.2e - 16$.

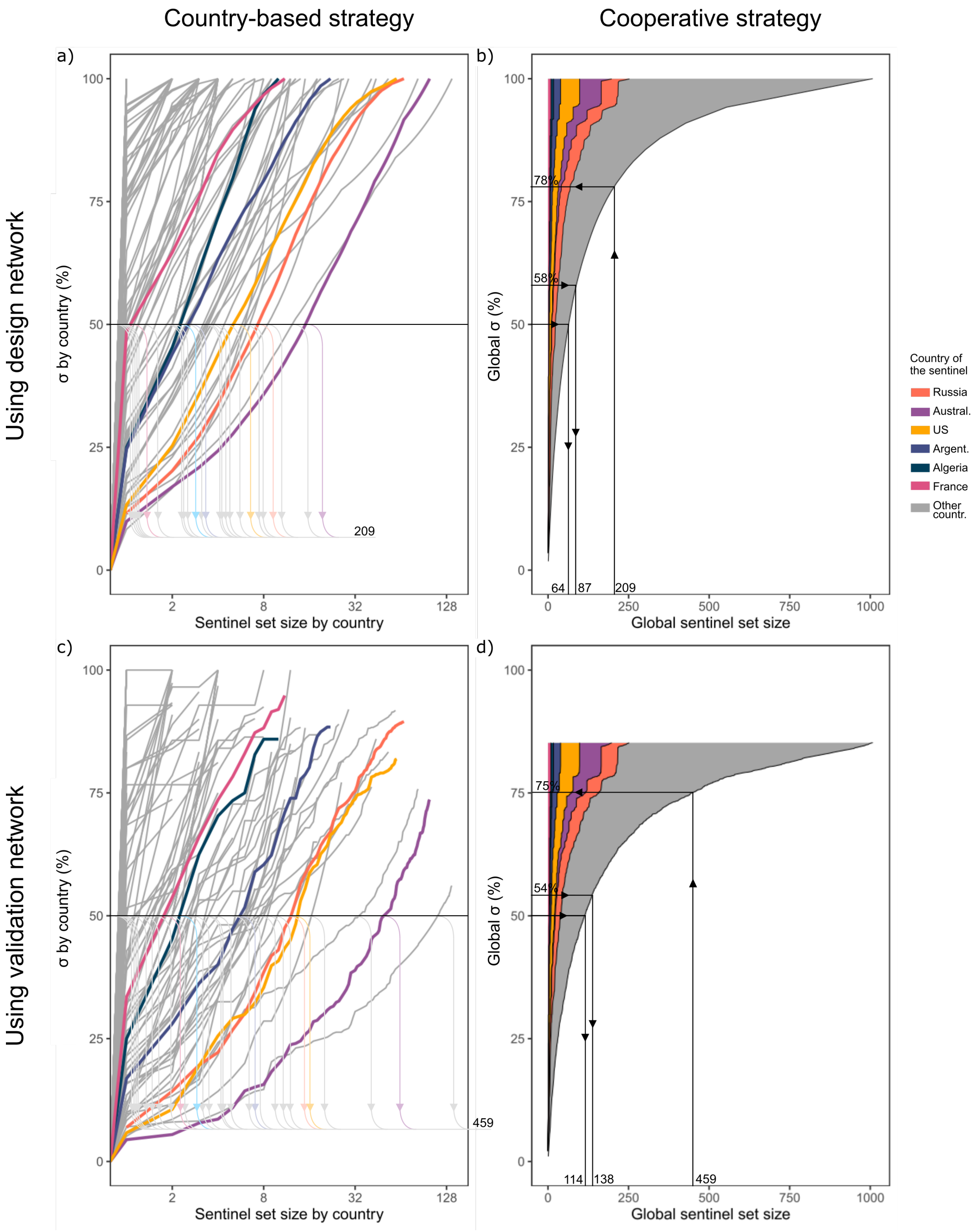


Figure SI2: Curves describing the relationship between increasing sentinel set size and relative aggregate coverage under the "Country-based" or "Cooperative" strategies (by columns) and using the design or validation networks (by rows). The first row is identical to Fig. 2 in the main text, except for panel b), where we highlight the fact that 87 sentinels are needed to achieve a target of 58%, corresponding to the effective aggregated coverage obtained with 209 sentinels in the "Country-based" scenario. In the second row, the performances of the same sentinels set are assessed using the validation network $\mathbf{C_V}$. In c) the "Country-based" strategy is reported; the number of sentinels needed to achieve a target of at least $\sigma = 50\%$ by country increases to 459 (Finland and Tunisia excluded). In d), the "Cooperative" strategy, where worldwide coverage of $\sigma = 50\%$ needs 114 sentinels, and 459 sentinels would allow a global coverage of $\sigma = 75\%$. A target of 54% (corresponding to the aggregated coverage obtained with 459 sentinels in the "country-based" scenario) needs 138 sentinels.

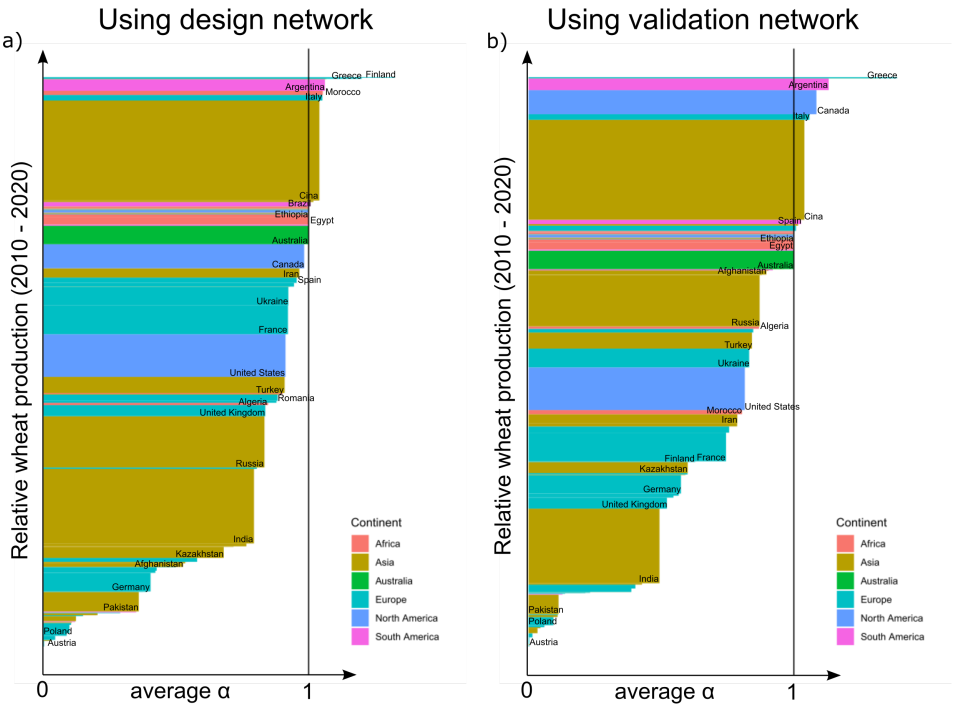


Figure SI3: Barplot of $\overline{\alpha}_i$ all over the world. Each country is represented by a rectangle where the base is proportional to $\overline{\alpha}$ and the height to wheat production in 2010 - 2020 [1]. In a) $\overline{\alpha}_i$ is calculated against the design network, in b) against the validation network.

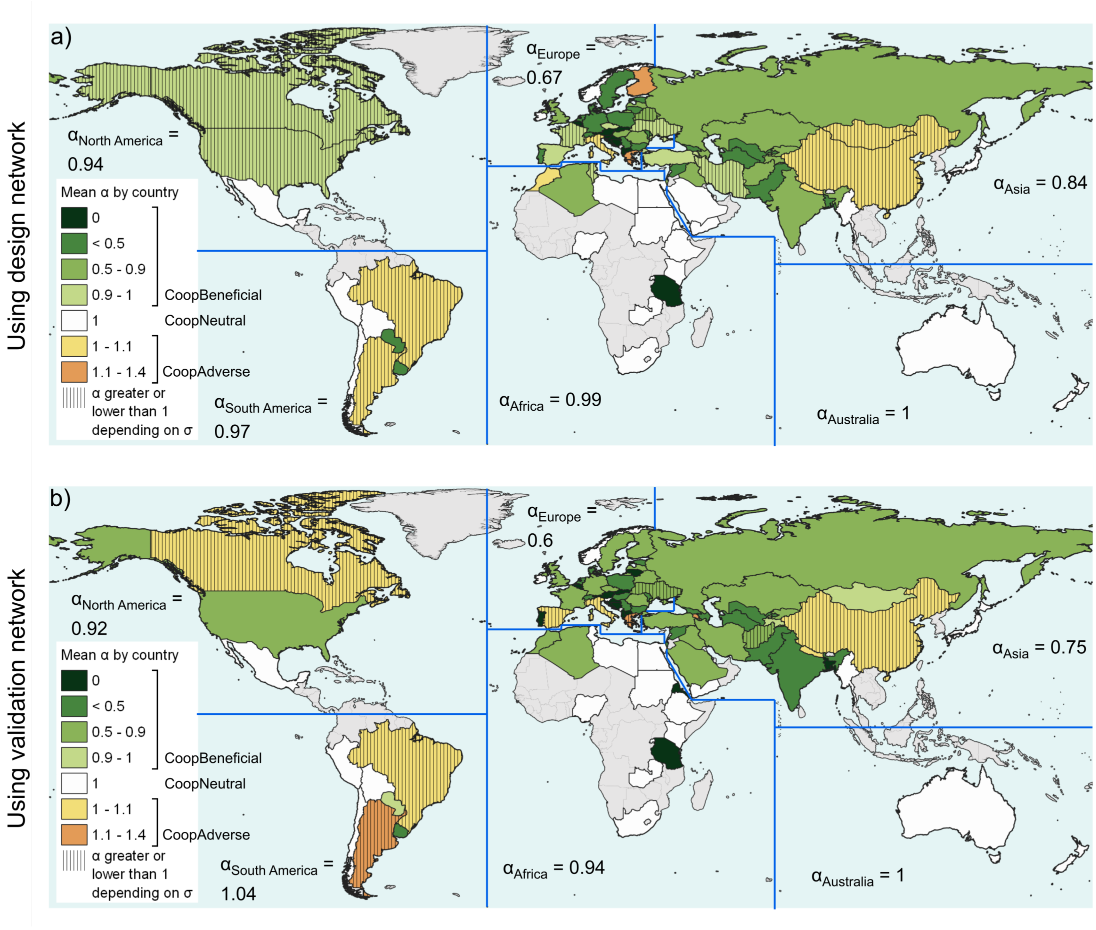


Figure SI4: Map of $\overline{\alpha}_i$ all over the world. On the top a) surveillance performances are assessed against the the design network, while on the bottom b) surveillance performances are assessed against the the validation network. Average values by continents, weighted by country wheat production 2010 - 2020 [1] are displayed.

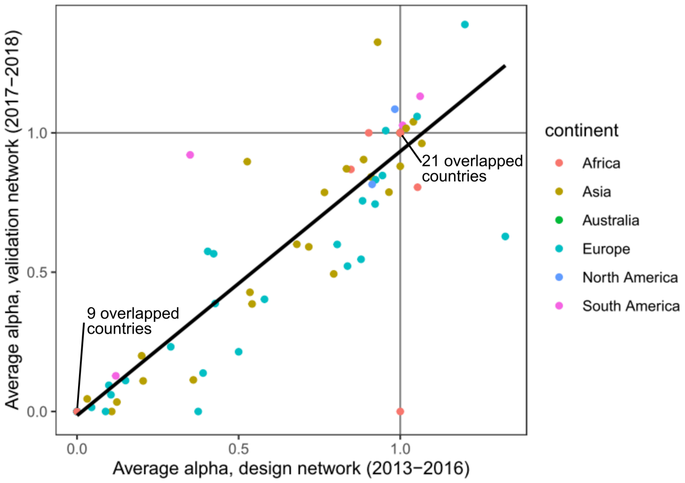


Figure SI5: Scatter plot representing the change in $\overline{\alpha}_i$ by country from design $C_D$ to validating networks $C_V$. The linear correlation coefficient among $\overline{\alpha_i}$ is 0.89, and the resulting p-value via the Pearson test is $< 2.2e - 16$. 77 out of 87 countries (i.e., 89% of the countries) keep qualitatively the same performances (i.e., $\overline{\alpha}_i$ is either $<$, $=$, or $> 1$ in both design $C_D$ and validation network $C_V$).
